# Separate, Separated and Together: the Transcriptional Program of the *Clostridium acetobutylicum- Clostridium ljungdahlii* syntrophy leading to interspecies cell fusion

**DOI:** 10.1101/2025.01.10.632433

**Authors:** Noah B. Willis, Eleftherios T. Papoutsakis

## Abstract

Syntrophic cocultures (hitherto assumed to be commensalistic) of *Clostridium acetobutylicum* and *Clostridium ljungdahlii*, whereby CO_2_ and H_2_ produced by the former feeds the latter, result in interspecies cell fusion involving large scale exchange of protein, RNA and DNA between the two organisms. Although mammalian cell fusion is mechanistically dissected, the mechanism for such microbial-cell fusions is unknown. To start exploring this mechanism, we used RNA sequencing to identify genes differentially expressed in this coculture using two types of comparisons. One type compared coculture to the two monocultures, capturing the combined impact of interactions through soluble signals in the medium and through direct cell-to-cell interactions. The second type compared membrane separated versus unseparated cocultures, isolating the impact of interspecies physical contact. While we could not firmly identify specific genes that might drive cell fusion, consistent with our hypothesized model for this interspecies microbial cell fusion, we observed differential regulation of genes involved in *C. ljungdahlii’s* autotrophic Wood-Ljungdahl-Pathway metabolism and genes of the motility machinery. Unexpectedly, we also identified differential regulation of biosynthetic genes of several amino acids, and notably of arginine and histidine. We verified that they are produced by *C. acetobutylicum* and are metabolized by *C. ljungdahlii* to its growth advantage. These and other findings, and notably upregulation of *C. acetobutylicum* ribosomal-protein genes, paint a more complex syntrophic picture and suggest a mutualistic relationship, whereby beyond CO_2_ and H_2_, *C. acetobutylicum* feeds *C. ljungdahlii* with growth boosting amino acids, while benefiting from the H_2_ utilization by *C. ljungdahlii*.

**IMPORTANCE:** The construction and study of synthetic microbial cocultures is a growing research area due to the untapped potential of defined multi-species industrial bioprocesses and the utility of defined cocultures for generating insight into complex, undefined, natural microbial consortia. Our previous work showed that coculturing *C. acetobutylicum* and *C. ljungdahlii* leads to a unique metabolic phenotype (production of isopropanol) and heterologous cell fusion events. Here, we used RNAseq to explore genes involved in and impacted by these fusions. First, we compared gene expression in coculture to each individual monoculture. Second, we utilized a transwell system to compare gene expression in mixed cocultures to cocultures with both species physically separated by a permeable membrane, isolating the impact of interspecies “touching” on the transcriptome. This study deepens our mechanistic understanding of the *C. acetobutylicum-C. ljungdahlii* coculture phenotype, laying the groundwork for reverse genetic studies of heterologous cell fusion in *Clostridium* cocultures.

## INTRODUCTION

Many of the diverse functions performed in the biosphere rely on microbial species working together. For example, microbial communities improve their ability to survive and thrive through cross-feeding of nutrients, secretion of defense agents, and breakdown of inhibiting or toxic compounds (1). Syntrophic interactions may explain (2) the existence of many so-called “unculturable” microbes, microbial species whose DNA can be detected and sequenced from environmental samples, but which cannot be isolated and cultivated individually in the laboratory. The expanded metabolic capability and improved efficiency of microbial communities has sparked considerable interest in rational design of cocultures for biotechnological applications. Recent research has focused on developing defined cocultures of metabolically complementary organisms for improved performance and expanded capabilities with minimal genetic engineering (3, 4). A growing number of such studies have utilized one or more organisms from the genus *Clostridium* (5, 6) to take advantage of their diverse metabolic capabilities (7). Direct cell-to-cell contact appears to play a central role in some coculture systems. Charubin et. al (8) demonstrated that conversion of glucose to alcohols and CO_2_ recycling was maximized in a coculture of the solventogen *Clostridium acetobutylicum* with the acetogen *Clostridium ljungdahlii* that was allowed to physically interact compared to cocultures in which the two species were separated by a permeable membrane. Pande et. al cocultured bacterial strains engineered to overproduce a certain amino acid alongside the corresponding auxotroph and showed similar results; the auxotrophic strain could obtain the amino acids it needed when mixed with the overproducing strain, but it could not survive when the two strains were separated by a permeable membrane (9). Observing the formation of long, tube-like structures between the overproducers and the auxotrophs in the non-separated condition (but not the membrane separated controls), they hypothesized that direct cytoplasmic exchange of amino acids via these “nanotubes” enabled the auxotrophs to survive. Other forms of cell-to-cell interactions via nanotubes structures have been well documented (10–12). There have also been reported examples of cytoplasmic content exchange without evidence of nanotube formation (2, 13). In our lab, metabolic studies, flow cytometry and confocal microscopy suggested that *C. acetobutylicum* and *C. ljungdahlii* can directly exchange metabolites (8), fluorescently labelled proteins & RNA (2), and more recently DNA (14). The DNA exchange events observed between *C. acetobutylicum* and *C. ljungdahlii* bear some resemblance to the DNA-exchange events facilitated by Type VII secretion systems in mycobacteria (15). Like mycobacteria, which require direct cell to cell contact to exchange DNA, the protein, RNA, and DNA exchange events were abolished when *C. acetobutylicum* and *C. ljungdahlii* were separated by a permeable membrane. Examination of *C. acetobutylicum*-*C. ljungdahlii* cocultures with scanning electron microscopy demonstrated that the two cell types directly fuse at their poles. Using qPCR, Charubin et. al showed that upregulation of several well characterized solventogenic genes correlate with the improved solvent (alcohol) production (8). However, the genetic program and corresponding proteins that induce and facilitate the observed cross-species membrane fusion events remain unexplored. Understanding these novel events would advance fundamental knowledge regarding the impact of cytoplasmic exchange events on individual microbes and on the communities they form, likely inspiring novel biotechnological applications.

Although multi-species transcriptomic studies have been widely reported, many of these have focused on host-pathogen interactions; transcriptomic studies on interspecies syntrophic interactions are less common (16). To our knowledge, only one such study has been reported for *Clostridium* cocultures, a pairing of *C. autoethanogenum* and *C. kluyveri* (17), which reported gene expression results for *C. autoethanogenum* only. Here, we used RNAseq to dissect the *C. acetobutylicum* - *C. ljungdahlii* coculture employing two different studies: compare gene expression of coculture against the individual monocultures (Type I study), and also compare cocultures separated by a membrane allowing metabolite exchange but no cell-to-cell contact versus unseparated cocultures (Type II study). The goal of the Type I study was to quantify how the transcriptomes of both *C. acetobutylicum* and *C. ljungdahlii* change in the coculture condition compared to typical monoculture growth conditions for each species. The goal of the Type II study was to isolate and quantify transcriptomic changes in *C. acetobutylicum* and *C. ljungdahlii,* which are specifically induced by direct cell-to-cell contact between the two species. Based on the analogous, well-characterized process in mammalian cells, we hypothesize that the cellular fusion events in our system involve dedicated proteins for chemotaxis and migration, membrane adhesion, and membrane fusion. Because cell surfaces carry a charge, membrane fusion events cannot happen spontaneously; energy-coupled protein machinery is required in order for cells to sense one another’s presence, seek each other out, initiate contact, and perform membrane fusion. Specifically, dedicated “fusogen” proteins are required to overcome the energetic barriers associated with the three key steps of the cell membrane fusion process: dehydration, hemifusion, and pore opening (18). In Gram-positive bacteria, such as both *C. acetobutylicum* and *C. ljungdahlii,* the additional barrier of the prokaryotic cell wall must also be overcome. Thus, to identify putative bacterial cell fusion enabling proteins, we specifically examined our coculture RNAseq results for differentially regulated genes known or annotated to be involved in chemotactic sensing, cellular motility, remodeling of the bacterial cell wall, or binding to the cell membrane.

## RESULTS AND DISCUSSION

### Experimental design: bottle and transwell experiments for Type I & II RNAseq comparisons, and global summary of differential gene expression from the two comparisons

For the first set of RNAseq comparisons/experiments (***Type I***), *C. acetobutylicum* and *C. ljungdahlii* cocultures were carried out in static screw cap bottles with loosened lids inside an anaerobic chamber (atmosphere composed of 85% N_2_, 10% CO_2_, and 5% H_2_) using the complex (5 g/L yeast extract) Turbo CGM growth medium in which interspecies fusion events were originally observed (8). The screw cap bottle lids were loosened to allow accumulated gas to escape (gas accumulation in pressurized glucose fermentations containing *C. acetobutylicum* can lead to explosions). *C. acetobutylicum* monoculture controls were grown in identical conditions to the cocultures. *C. ljungdahlii* monoculture controls were grown in pressurized serum bottles and mixed on a rotary shaker at 90 rpm, instead of static de-pressurized screw cap bottles, so that the bottles could be charged with 20 psig of an H_2_/CO_2_ (80/20) gas mixture to simulate the CO_2_ and H_2_ produced by *C. acetobutylicum* in the coculture condition. *C. ljungdahlii* monocultures were grown in the same medium as the coculture and *C. acetobutylicum* monoculture controls except the medium used in the *C. ljungdahlii* monoculture controls did not contain any glucose. Since *C. ljungdahlii* cannot use glucose, the osmotic pressure of high glucose concentrations can slow *C. ljungdahlii growth*. In coculture, glucose concentrations are reduced by its utilization by *C. acetobutylicum*, thus reducing *C. ljungdahlii* inhibition.

The goal of the Type I comparisons was to assess how the interaction of the two species in the coculture affects overall gene expression patterns of each species compared to their respective monocultures. This is the typical type of published comparisons of coculture systems, as direct cell-to-cell contact effects had not been anticipated until recently as discussed. RNA was isolated from the coculture and the monocultures at 4, 11, and 22 hours for Type I comparisons (Fig. 1A). The 4 hour timepoint was chosen because our previous work has shown that cytoplasmic material exchange between the two organisms is the most active at this time (2). The 11 and 22 hour timepoints were chosen because they are representative of *C. acetobutylicum*’s acidogenic phase and its transition from the acidogenic phase to solventogenic phase, respectively. Three biological replicates were prepared for the Type I comparisons; growth, culture pH, and metabolite data from these cultures are summarized in Figs. 1B, C, & E. The population ratio of the two species in the coculture at each timepoint was estimated based on the number of aligned mRNA reads from each organism at each timepoint (Fig. 1D). As shown in Fig. 1C, pH was manually adjusted from below 5.0 to approximately 5.3 at 11 hours in all Type I cultures to support the transition to solventogenesis.

**Figure 1:**
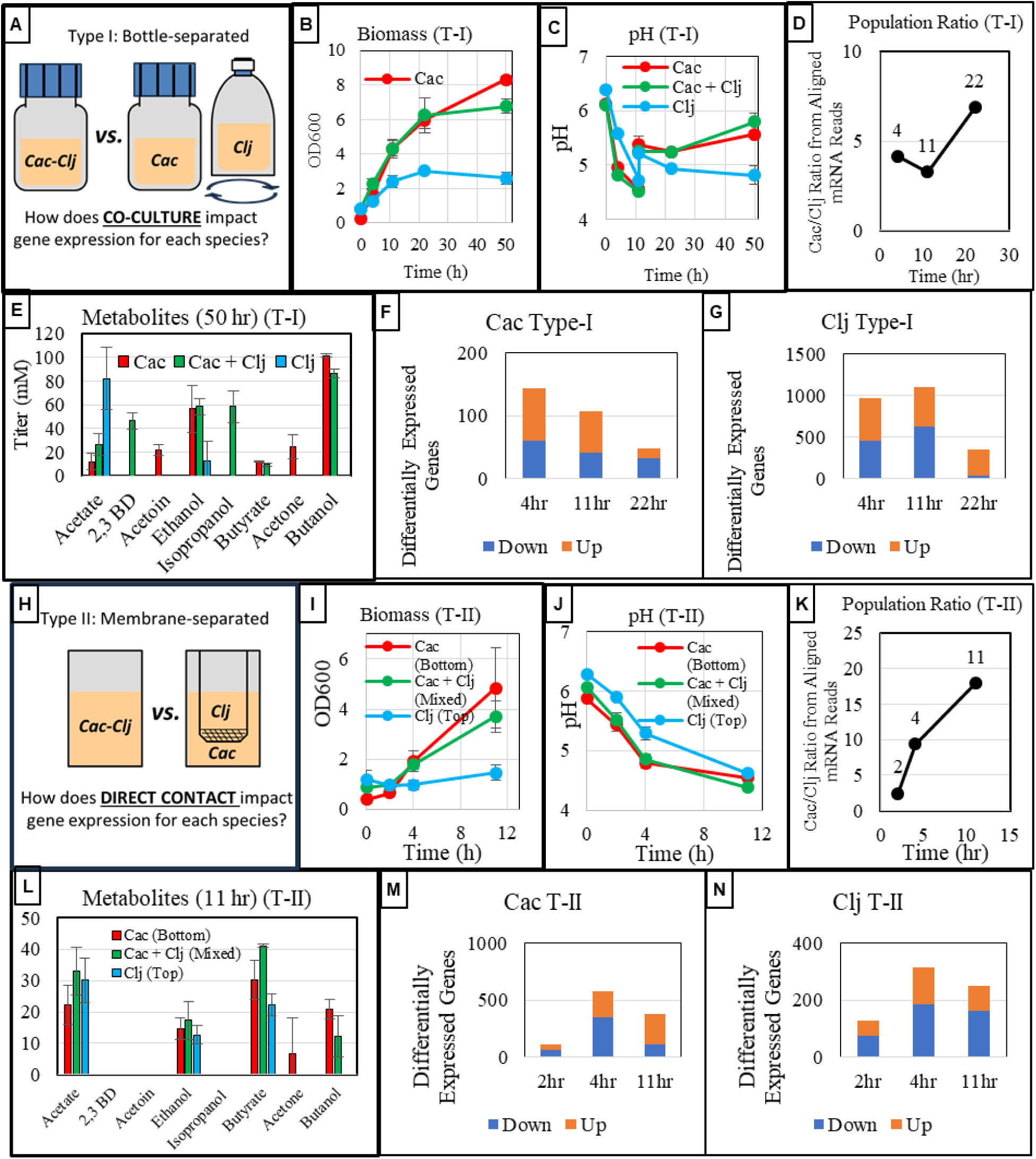
Summary of experimental design and outcomes. Three biological replicates were performed for both Type I and Type II cultures. A) Experimental setup for Type I experiments/comparisons. B) Biomass (OD_600_) profile for Type I cultures. C) pH profile for Type I cultures. D) C. acetobutylicum- C. ljungdahlii (Cac-Clj) population ratio based on aligned mRNA reads for Type I cultures. E) Metabolite titers after 50 hrs for Type I cultures. F) Differentially expressed genes in C. acetobutylicum in the Type I experiments. G) Differentially expressed genes in C. ljungdahlii in Type I experiments. H) Experimental setup for Type II experiments/comparisons, I) Biomass (OD_600_) profiles for Type II cultures. J) pH profiles for Type II cultures. K) C. acetobutylicum- C. ljungdahlii (Cac-Clj) population ratio based on aligned mRNA reads for Type II cultures. L) metabolite titers after 11 hrs for Type II cultures. M) Differentially expressed genes in C. acetobutylicum in the Type II experiments. N) Differentially expressed genes in C. ljungdahlii in the Type II experiments.

For the second set of RNAseq comparisons (***Type II***), using the same glucose containing medium as in the Type I coculture experiments, *C. acetobutylicum - C.ljungdahlii* cocultures were set up in transwells (Fig. 1H) with and without a separating 0.4-µm pore polycarbonate membrane. The membrane allows transport of small and large molecules between the two compartments, but it prevents direct contact between the cells of the two species. Due to the nature of the well plate, the transwell cultures could not be pressurized, so they were grown in the same anaerobic incubator and under the same atmosphere as described for the screw cap bottles in the Type I experiments. This transwell-system setup has previously been successfully used for this biological system (8) to demonstrate the importance of direct contact between the two species. *The goal of these Type II comparisons was to specifically examine how direct interspecies cell-to-cell contact impacts gene expression in coculture, while controlling for other forms of interspecies interactions that can occur through the liquid media, including gas exchange, diffusion of soluble metabolites, nutrient competition, and quorum sensing.* RNA was isolated from a mixed *C. acetobutylicum - C. ljungdahlii* coculture at 2, 4, and 11 hours and compared to RNA extracted from the top (*C. ljungdahlii* cells) and bottom (*C. acetobutylicum* cells) of the membrane separated cocultures (Fig. 1H). For the membrane separated cocultures, *C. ljungdahlii* was grown on the top compartment so that it could easily capture gases released by *C. acetobutylicum* from the bottom compartment. The 2 hour timepoint was chosen to obtain a snapshot of gene expression patterns before significant direct *C. acetobutylicum* to *C. ljungdahlii* cell interactions, leading to heterologous cell fusion, begin (8). The 4 and 11 hour timepoints were maintained in order to allow comparisons to the Type I coculture and monoculture experiments described above. Three biological replicates were carried out. Growth, culture pH, population ratio, and metabolite data from these cultures are summarized in Figs. 1I, J, K and L. pH was not manually adjusted in the Type II experiments because no timepoints after 11 hours were studied. We have consistently shown in previous work (8) and in the Type I experiments that both cocultures and monocultures are active at a pH below 5.0; manual pH adjustment is only beneficial for supporting robust growth after 11 hours. Biomass accumulation in the mixed transwell coculture at 11 hours in the Type II transwell system experiment (OD_600_ of 3.71; Fig. 1I) was similar to biomass accumulation in the coculture at 11 hours in the Type I bottle system experiment (OD_600_ of 4.32; Fig. 1B), showing that active growth was successfully maintained up until 11 hours despite the low pH, at which point all remaining cells were harvested for RNA extraction. There was no detectable isopropanol production in the transwell coculture at 11 hours (Fig. 1L), but this was not unexpected since acetone production by *C. acetobutylicum* (and subsequent conversion to isopropanol by *C. ljungdahlii*) is minimal or nonexistent in the acidogenic phase, as seen from the Type I coculture metabolite profile (Fig. S1A). Hereafter, when discussing gene expression, genes expressed “higher”, “lower” or alternatively “upregulated”, “downregulated” will always refer to the coculture relative to monoculture (for Type I comparisons) and unseparated coculture relative to separated coculture (for Type II comparisons). Complete summaries of both the differential and absolute gene expression results for *C. acetobutylicum* and *C. ljungdahlii* for both the Type I and II RNAseq experiments are available in the supplementary material (Tables S1-S4).

In Type I experiments, the *C. acetobutylicum* presence impacted the *C. ljungdahlii* transcriptome far more than the *C. ljungdahlii* presence impacted the *C. acetobutylicum* transcriptome (Fig. 1F, G). At the 4, 11, and 22-hour fermentation timepoints sampled, 21%, 25%, and 8% of *C. ljungdahlii* genes were differentially expressed compared to the *C. ljungdahlii* monoculture, respectively. In contrast, less than 5% (at any timepoint) of *C. acetobutylicum* genes were differentially expressed compared to the *C. acetobutylicum* monoculture. These findings are consistent with modelling work of this syntrophic system suggesting that, in order to persist in the coculture with the much faster growing *C. acetobutylicum*, the concomitant accumulation of primary metabolites, and the fast production of CO_2_ and H_2_, *C. ljungdahlii* metabolism is more profoundly impacted in this syntrophy (19). These large *C. ljungdahlii* transcriptional changes reflect its survival response to the fast production of CO_2_ and H_2_ by *C. acetobutylicum* and the accumulation of primary metabolites, combined with the depletion of fructose, and the competition for amino acids from the yeast extracts in the medium. While the CO_2_ and H_2_ support *C. ljungdahlii* growth and survival as fructose is being depleted, inhibitory metabolites (such as butyrate and butanol) and competition for yeast-extract amino acids from the dominating *C. acetobutylicum* drives *C. ljungdahlii* to adopt a survival strategy. With some success as shown in Fig. 1D: it succeeds in enriching in the population from hour 4 to 11, but cannot sustain the fight to hour 22, possibly due to accumulation of *C. acetobutylicum* toxic metabolites.

In Type II experiments, a slightly larger percentage of *C. acetobutylicum* genes was differentially expressed compared to *C. ljungdahlii* (Fig. 1M, & N). At the 2, 4, and 11-hour timepoints, 3%, 15%, and 10% of *C. acetobutylicum* genes were differentially expressed, compared to 3%, 7%, and 6% of *C. ljungdahlii* genes at the same timepoints. Previous flow cytometry data (2) suggest that *C. acetobutylicum*-*C. ljungdahlii* cytoplasmic exchange increases substantially between 1 and 4.5 hours of coculture. As the Type II comparisons assess the impact of direct cell-to-cell contact (including interspecies cell fusion) vs. indirect communication via soluble, transwell-membrane permeable signals, these results are consistent with our previous work, as there is a very large increase in the number of differentially expressed genes from 2 to 4 hours in both species.

### Overview of Genes Strongly Up-Regulated by Direct Interspecies Contact (Type II comparisons)

As a first pass to identify protein candidates that may be engaged in interspecies cell fusion, we examined the top twenty (Fig. 2) and top 50 (Tables S5, S6) most upregulated genes in both species in the membrane-separated (Type II) experiments. We examined these genes aiming to identify genes/proteins that characterize the response of the cells to direct interspecies contact, including specifically those involved in facilitating direct interspecies contact. We include the “blue” expression abundance RPKM (reads per kilobase per million mapped reads) plots to ascertain that the genes discussed are expressed at levels commensurate with their expected function. We focused on the 4-hour timepoint because, as discussed, this seems to be an early hotspot for interspecies fusion based on our prior work (2).

**Figure 2:**
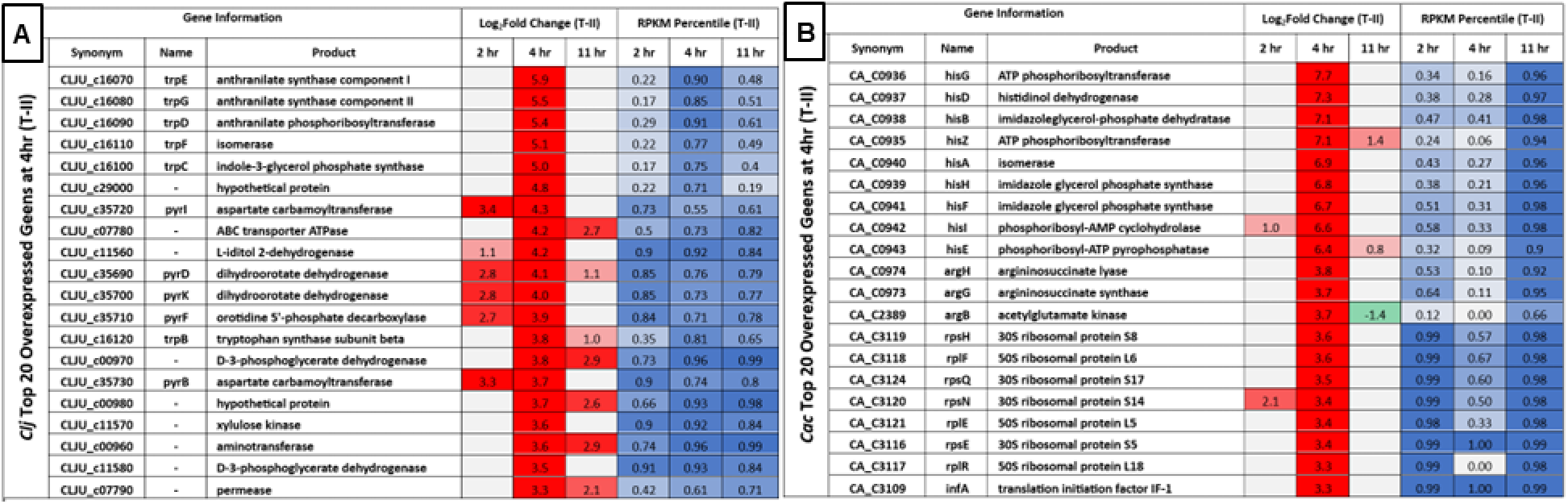
Top 20 differentially up-regulated genes at the 4 hr timepoint sorted by log2fold change in both C. acetobutylicum and C. ljungdahlii in the Type II experiments. Red (upregulated) and green (downregulated) cells in the log_2_fold change heatmap columns indicate genes differentially regulated with statistical significance between conditions at each timepoint. Blue cells in the RPKM (reads per kilobase per million mapped reads) heatmap columns represent the percentile of transcript abundance for each gene relative to all of the other genes in the genome at each timepoint.

Many of the strongly upregulated (Fig. 2, Tables S5 & S6), and largely highly expressed, genes in both *C. acetobutylicum* and *C. ljungdahlii* at the 4 hour timepoint in the Type II experiments were annotated as genes involved in established pathways for amino acid biosynthesis, protein translation, and pyrimidine biosynthesis. Strong enrichment of ribosomal proteins (in *C. acetobutylicum*) and pyrimidine biosynthesis (rRNA in *C. ljungdahlii*) among highly upregulated genes in both organisms suggests that direct cell-to-cell contact stimulates cell growth of both organisms, as suggested by prior modelling work of this coculture system (19). Upregulation of ribosome and pyrimidine biosynthesis (*pyr* genes) attends to the needs of the cell for faster growth. But what drives the need for faster growth of the two organisms under coculture? Likely reasons are detailed in subsequent sections and include novel findings of physiological significance.

While interesting and consistent with the observed coculture phenotype, none of the highly upregulated genes discussed above appear to be obvious candidates for novel prokaryotic “fusogen” genes which directly carry out the process of cell wall and membrane fusion. Two *C. ljungdahlii* genes of potential interest among the top 20 differentially upregulated genes (Fig. 2) are the two contiguous genes, CLJU_c07780 (putative ABC transport system ATP-binding protein) and CLJU_c07790 (putative ABC transport system permease). Both genes are strongly upregulated at the 4 and 11 hour timepoints. In the Pfam database, the permease belongs to the FtsX-like permease family, a group of ATP-requiring transporter proteins used to move lipids across cell membranes (20). The canonical FtsX and its associated ATP-binding protein FtsE are established as widely-conserved important regulators of cell wall peptidoglycan hydrolases (21), so it is possible these *C. ljungdahlii* genes may play a role in the cell wall and membrane remodeling process during cell fusion.

In *C. acetobutylicum,* virtually all of the top 50 upregulated genes at 4 hours are well annotated and unlikely to be part of the machinery involved in interspecies contact and material exchange (Fig. 2, Table S5). Among the few poorly annotated genes upregulated in *C. acetobutylicum* at the 4-hour timepoint in the Type II experiment, we identified that three genes (CA_C0042, 47-48) from a putative Type VII secretion system were upregulated at a low level in response to direct contact with *C. ljungdahlii* (Table S7) and a fourth gene in the operon, CA_C0039, narrowly missed the cutoff for statistically significant upregulation (p = 0.09). Type VII secretion systems (originally known as the ESAT-6/WXG100 secretion system) have only been characterized and studied in *C. acetobutylicum* bioinformatically (22). However, extensive work has documented the Type VII systems in *Mycobacterium spp,* where they have a variety of contact dependent functions (15). One of these mycobacterial Type VII systems is believed to be responsible for the large scale intraspecies chromosomal DNA exchange and “mosaic” genomes in *Mycobacterium tuberculosis* and *Mycobacterium canetti* (23). The diverse functions of these secretion systems are still an active area of study in *Mycobacterium* and other microbes, including recently in the model Gram positive *Bacillus subtilis* (24). *C. acetobutylicum* has three putative operons coding for Type VII systems, though upregulation at 4 hours was only observed for genes of the CA_C0035-0049 operon. Based on BLAST analysis, *C. ljungdahlii* does not contain any putative Type VII genes. However, Type VII systems have been shown to be polar, requiring the secretion system machinery only in the recipient cells (25), here, hypothetically, *C. acetobutylicum*.

Although we hypothesized that there would be strong upregulation of several genes that might be involved in heterologous cell fusion in response to interspecies contact, it is possible that many such genes would only be strongly upregulated in individual cells that are about to undergo or are actively undergoing cell fusion. We have shown that at any time a maximum of approximately 14% of cells are involved cell fusion events (2). Thus, the bulk isolation of RNA from the entire population, such as performed in our study, likely leads to “dilution” of RNA from actively fusing cells by RNA from the balance of the cell population. Based on these results, quantification of the transcriptome of actively fusing hybrid cells may require a combination of fluorescently labelled *C. acetobutylicum* and *C. ljungdahlii* strains for cell sorting with a transcriptomic analysis on a sorted cell population (Sort-Seq).

### Arginine, histidine, and tryptophan biosynthesis genes and metabolism

As stated above, many of the pathways that were differentially regulated with statistical significance involve amino acid metabolism and transport, among which, three amino acids clearly stood out: arginine, histidine, and tryptophan. We thus analyzed supernatant samples from coculture and both monocultures to evaluate the profiles of these three amino acids over the course of the entire Type I experiment (Fig. 3). All amino acids in the starting medium derive from the yeast extract (5 grams per liter). The profiles of the three amino acids in monocultures suggest that *C. ljungdahlii* consumes all three amino acids available in the medium very fast and must then start their biosynthesis (given the null concentrations in the medium) within 4 hours upon inoculation for arginine and histidine, and from the very beginning for tryptophan. The *C. acetobutylicum* monoculture profiles suggest that *C. acetobutylicum* does not need to synthesize arginine or histidine until after 20 hours. Like *C. ljungdahlii*, tryptophan needs to be synthesized by *C. acetobutylicum* from the very beginning. The coculture amino-acid profiles, however, are virtually identical to the *C. ljungdahlii* profiles. This would suggest that *C. acetobutylicum,* as the major feeder of the coculture through glucose catabolism, must upregulate the biosynthesis of arginine or histidine acids within 4 hours, and this is indeed what the transcriptional data suggest. In this sense, *C. acetobutylicum,* does not only provide CO_2_ and H_2_ to *C. ljungdahlii* for its growth and survival, but also amino acids, which likely enable faster growth as detailed next.

**Figure 3:**
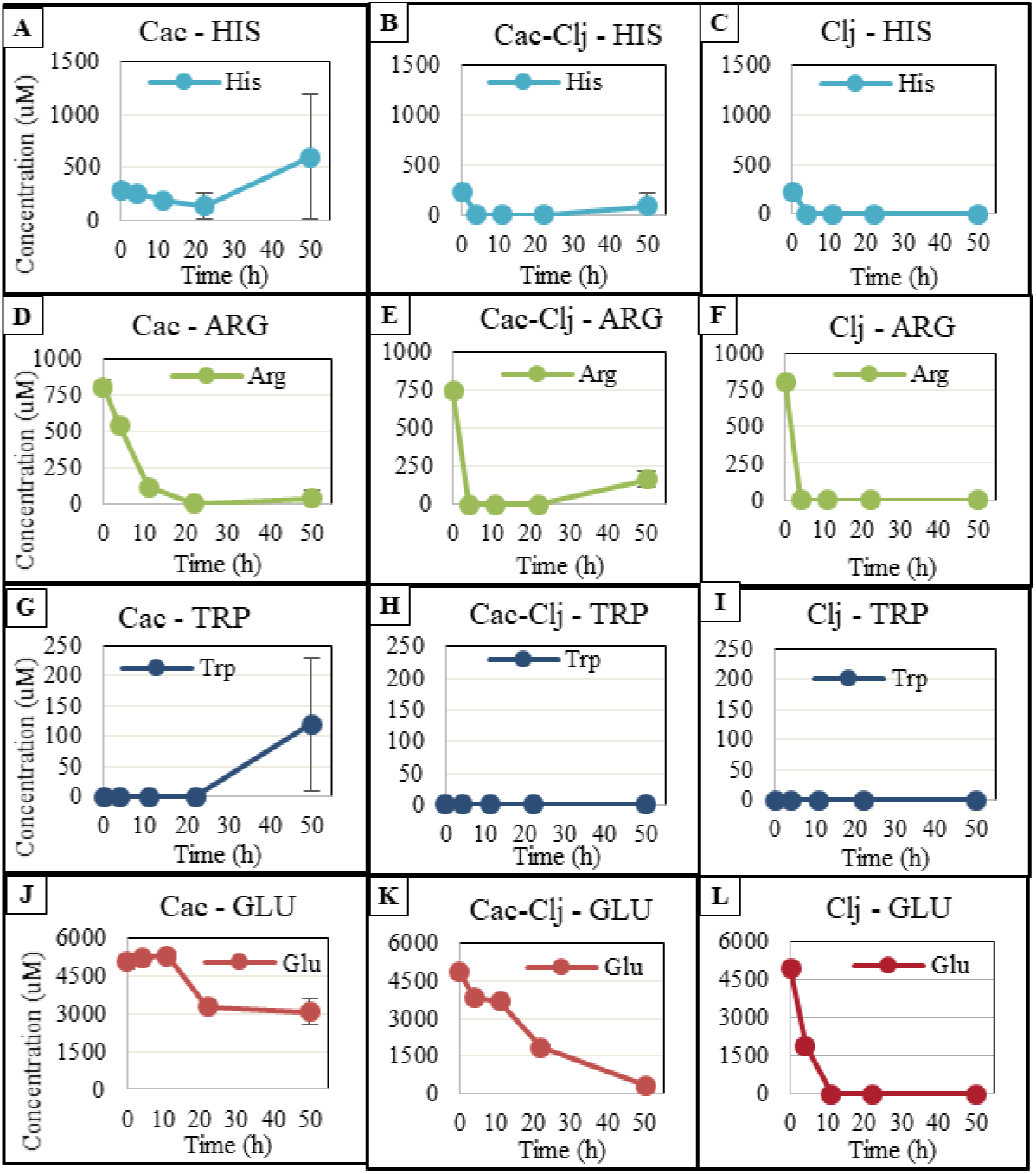
Time profiles of arginine, histidine, and tryptophan concentrations in the medium for *C. acetobutylicum* monoculture, *C. ljungdahlii* monoculture, and *C. acetobutylicum- C.* ljungdahlii cocultures for Type I experiments (N=3). ‘N’ represents the number of biological replicates used for analysis and error bars represent standard deviation of the three replicates at each timepoint.

For arginine biosynthesis in the Type I experiments in *C. acetobutylicum*, five of the central genes were strongly upregulated at 4 hours (CA_C0316, 0973-0974, 2388, 2391) and three of those genes were also upregulated at 11 hours (Fig. 4A). In the transwell experiments, eight of the central arginine biosynthesis genes were upregulated at 4 hours (Figure 4A). In *C. ljungdahlii* many of the central arginine biosynthesis genes are either up or down regulated in both the Type I and Type II experiments with no clear pattern (Fig. 4B). However, all three genes (CLJU_c09270, 09280, 09300) of the arginine deiminase catabolic pathway were upregulated at 4 hours in the Type I experiments and at 4 and 11 hours in the Type II experiments. From one mole arginine, the arginine deiminase pathway generates one mole each of ornithine, ATP, and CO_2_, and two moles ammonia. Under both heterotrophic and autotrophic growth conditions (26), arginine catabolism via the arginine deiminase pathway has been shown to increase the growth rate of *Clostridium autoethanogenum*, an acetogen virtually identical to *C. ljungdahlii. C. acetobutylicum* does not have homologs for the arginine deiminase pathway. These results from the Type II comparisons (differential expression due to direct cell contact), suggest that, via direct contact in the coculture system, *C. ljungdahlii* may utilize arginine produced by *C. acetobutylicum* for additional ATP production.

**Figure 4:**
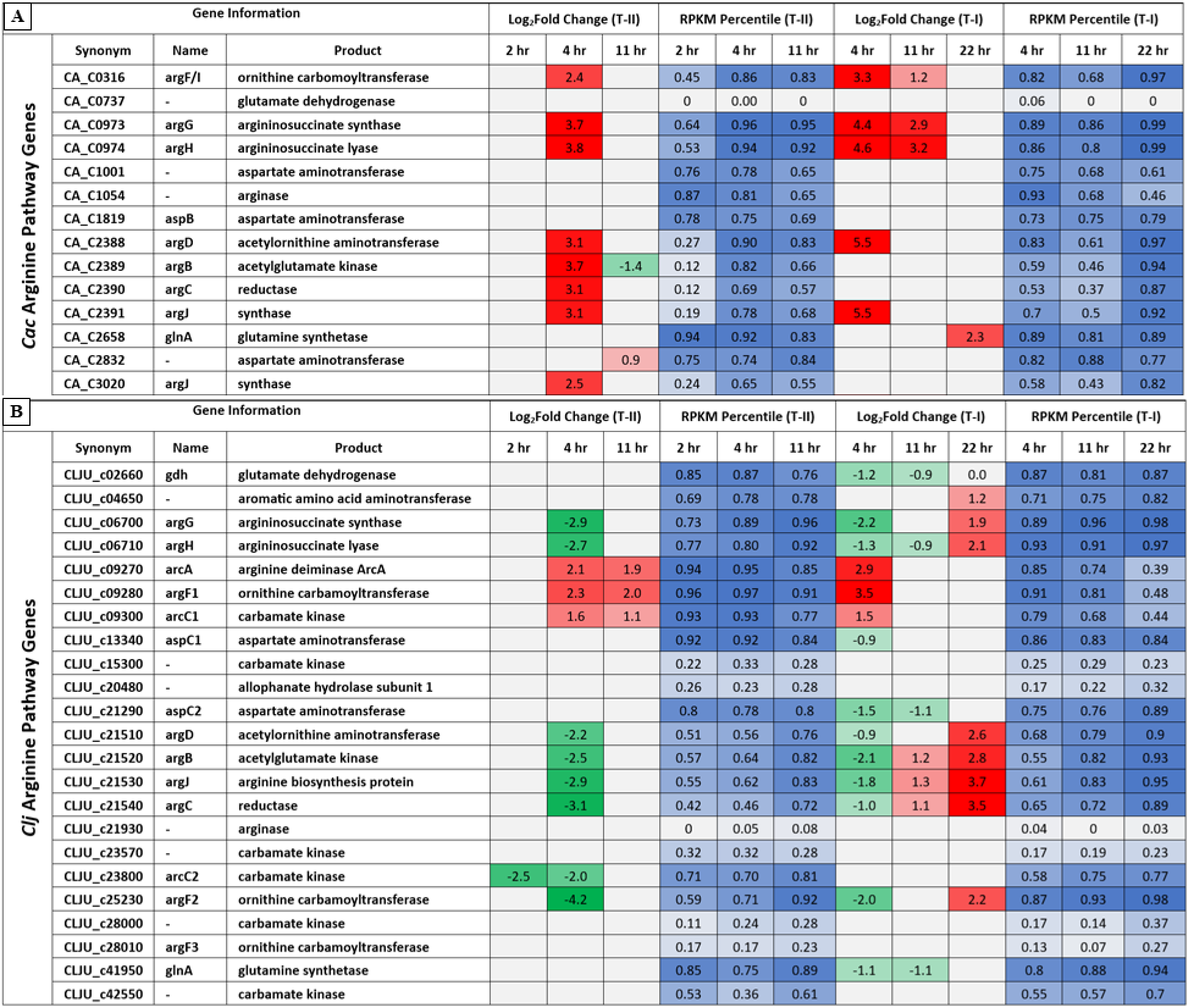
Heatmap of differential gene expression of arginine biosynthesis genes in Type I and Type II comparisons for A) C. acetobutylicum (Cac) and B) C. ljungdahlii (Clj). Red (upregulated) and green (downregulated) cells in the log2fold change heatmap columns indicate genes differentially regulated with statistical significance between conditions at each timepoint. Blue cells in the RPKM ((reads per kilobase per million mapped reads) heatmap columns represent the percentile of transcript abundance for each gene relative to all of the other genes in the genome at each timepoint.

The genes of the histidine biosynthesis pathway at 4 and 11 hours were among the most strongly upregulated in *C. acetobutylicum* in the Type I experiments (Fig. 5A). A similar, strong upregulation of the entire operon was observed at 4 hours in the Type II transwell experiments (Fig. 5). For *C. ljungdahlii*, many of the histidine biosynthesis genes were upregulated (Figure 5B). However, more interestingly, four genes involved in histidine degradation to glutamate (CLJU_c21460-21480, 21500) were strongly upregulated early in coculture. If the reactions catalyzed by the protein products of these genes are summed (EC#s 4.3.1.3, 4.2.1.49, 3.5.2.7, 2.1.2.5), the net reaction is the combination of histidine and a tetrahydrofolate (THF) group to produce ammonia, glutamate, and 5-formimidoyl-THF (Eq. 1). The *C. ljungdahlii* glutamate formiminotransferase (CLJU_c21500), is highly homologous to eukaryotic formimidoyltransferase-cyclodeaminases (which also carry the glutamate formiminotransferase activity), which catalyze the breakdown of 5-formimidoyl-THF into ammonia and 5,10-Methenyl-THF (EC 4.3.1.4). If we add this additional reaction (Eq. 2), the net reaction is degradation of histidine to ammonia, glutamate (GLU) and the key WLP eastern branch intermediate 5,10-Methenyl-THF (Eq. 3).

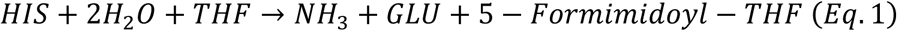

**Figure 5:**
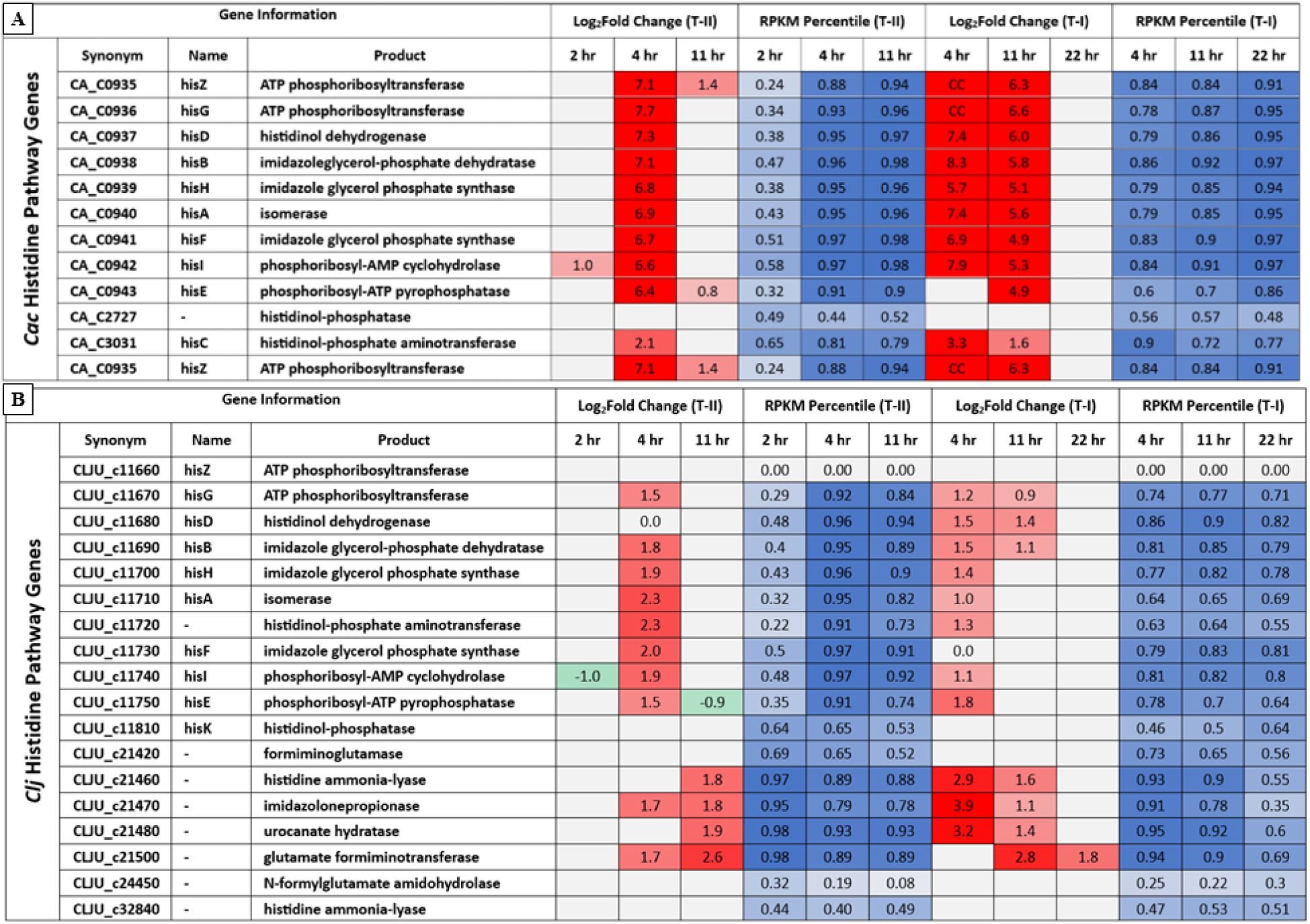
Heatmap of differential gene expression of histidine biosynthesis genes in Type I and Type II comparisons for A) C. acetobutylicum (Cac) and B) C. ljungdahlii (Clj). Red (upregulated) and green (downregulated) cells in the log2fold change heatmap columns indicate genes differentially regulated with statistical significance between conditions at each timepoint. Blue cells in the RPKM ((reads per kilobase per million mapped reads) heatmap columns represent the percentile of transcript abundance for each gene relative to all of the other genes in the genome at each timepoint.

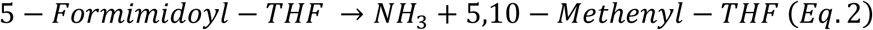

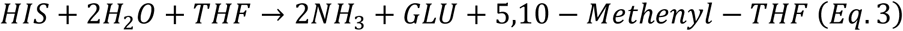

Normally, generation of 5,10-methenyl-THF from formate in the eastern branch of the WLP requires ATP. This ATP consuming step makes the overall WLP pathway ATP limited for growth on CO_2_, requiring the production of acetate via substrate-level phosphorylation from acetyl-CoA (the end product of the WLP) for significant ATP production to support cell growth (27). The ATP generated by the -complex mediated proton gradient (28) is not sufficient to support cell growth. Production of 5,10-methenyl-THF from histidine skips this step, increasing the energy efficiency of the WLP while generating one mole of glutamate and two moles of ammonia per mole histidine consumed. This hypothesized explanation for increased histidine synthesis by *C. acetobutylicum* and increased histidine utilization by *C. ljungdahlii* is further supported by the monoculture and coculture secretion profiles of glutamate (Fig. 3). In their respective monocultures, *C. acetobutylicum* utilizes less than half of the available glutamate whereas *C. ljungdahlii* quickly utilizes all of the available glutamate by the 11 hour timepoint. However, in coculture the rate of consumption is much slower than in the *C. ljungdahlii* monoculture, and there is still residual glutamate at the 50 hour timepoint, which could be explained by decreased glutamate requirement of *C. ljungdahlii* in coculture due to increased intracellular glutamate generation via histidine catabolism.

To test the suggested hypothesis that arginine and histidine increase *C. ljungdahlii’*s growth, we tested the autotrophic growth of *C. ljungdahlii* under three conditions: supplementing 5 g/L arginine (28 mM), 5 g/L histidine (32 mM), or 5 g/L each of both arginine and histidine. The basal medium to which these supplemental amino acids were added was the Turbo CGM medium made without glucose or fructose. A fourth control condition was included in which *C. ljungdahlii* was grown in Turbo CGM without sugars or any of the supplemental amino acids. We also measured the ammonia-formation kinetics since ammonia is a product of both the arginine deiminase pathway and the proposed pathway for histidine degradation. As expected, based on previous results from *C. autoethanogenum* (26), arginine addition significantly boosted the growth rate and total biomass accumulation of *C. ljungdahlii* (Fig. 6A). However, the total production of acetate, and thus CO_2_ fixation (as CO_2_ is the main carbon source of these cultures with and especially without the amino acid supplementation), decreased substantially (33 mM net production of acetate with arginine; 88 mM net production of acetate without) (Fig. 6B). Although counterintuitive, this makes sense metabolically. The arginine deiminase pathway is completely orthogonal to the WLP. With easily available ATP from arginine catabolism, *C. ljungdahlii* does not need to rely as heavily on CO_2_ utilization via the WLP, as it produces relatively little ATP from it. In contrast, the cultures growing on histidine made more acetate (142 mM), meaning they fixed substantially more CO_2_. This makes sense because, based on the proposed pathway to 5,10-methenyl-THF, histidine catabolism is synergistic with the WLP. Even the cultures fed both arginine and histidine produced less acetate (and thus fixed less CO_2_) than the cultures fed only histidine (Fig. 6B). Ammonia formation was consistent with the observed growth patterns and proposed amino acid catabolism pathways (Fig. 6C). These results suggest that both arginine and histidine can substantially improve *C. ljungdahlii* growth, explaining why *C. ljungdahlii* may seek to obtain these amino acids in coculture with *C. acetobutylicum.* However, of the two, histidine was superior both for increasing growth and CO_2_ fixation by *C. ljungdahlii*.

**Figure 6:**
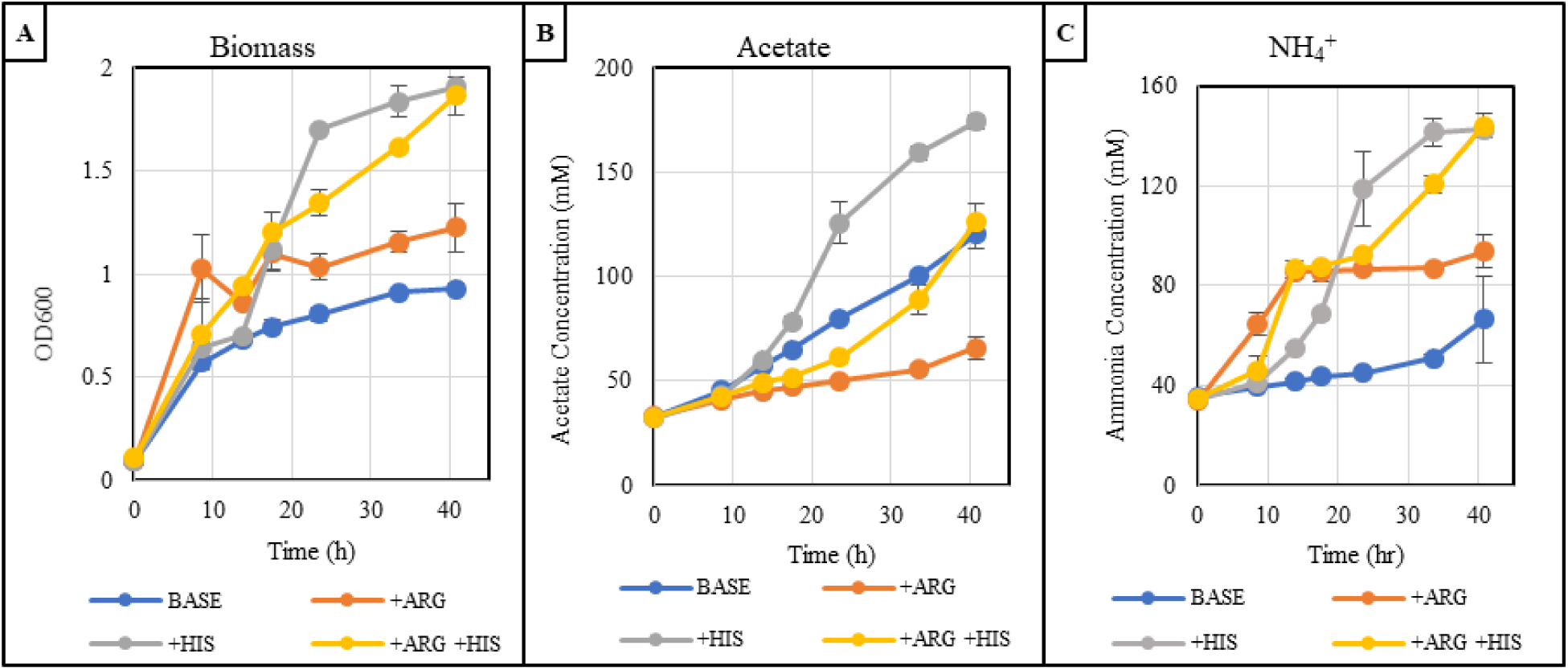
Temporal profiles of (A) OD_600_ (biomass), (B) acetate, and (C) ammonia for autotrophic C. ljungdahlii monocultures supplemented with either 5 g/L arginine, 5 g/L histidine, or 5 g/L of both arginine and histidine compared to control (no supplementation) (N=2). ‘N’ represents the number of biological replicates used for analysis and error bars represent standard deviation of the two replicates at each timepoint. Ammonium present at the beginning of the experiment is from the ammonium sulfate provided in the media.

According to the KEGG database, *C. acetobutylicum* does not have the genes required for histidine conversion to glutamate. Even if *C. acetobutylicum* did have the genes, the monoculture was clearly not glutamate limited, and, since it lacks the WLP, *C. acetobutylicum* would not derive any obvious benefit from generating additional 5,10-methenyl-THF. The same study that identified the growth enhancing mechanism of arginine via the arginine deiminase pathway in *C. autoethanogenum* also identified histidine as a potentially growth enhancing amino acid via “shadow price analysis”, but they did not propose a mechanism (26). We hypothesize that the strong pattern of upregulation of histidine biosynthesis genes observed in our experiments for both organisms, but especially in *C. acetobutylicum*, is explained by the cross-feeding of histidine from *C. acetobutylicum* to *C. ljungdahlii* which *C. ljungdahlii* catabolizes to increase the energy efficiency of CO_2_ fixation via the WLP.

In contrast to differential expression of histidine and arginine genes, which largely happens early in coculture during the growth phase, genes related to tryptophan biosynthesis were strongly upregulated at later timepoints. In *C. acetobutylicum* Type I experiments, seven of the key tryptophan biosynthesis genes were strongly upregulated at 22 hrs (CA_C3157-3163) (Table S8). In *C. ljungdahlii* the seven homologous genes were also strongly upregulated at 22 hr (CLJU_c16070-16130) (Table S9). These genes were also upregulated strongly in *C. ljungdahlii* in the transwell experiments (Type II). We cannot yet interpret how these findings relate to the tryptophan profile in the culture medium (Fig. 3)

### Does the differential expression of ribosomal proteins support the notion that the coculture benefits the growth of *C. acetobutylicum?*

As stated above, of the top 50 most differentially expressed genes in *C. acetobutylicum* at the 4 hour timepoint in the Type II experiments, nineteen genes were annotated as ribosomal proteins (Fig. 2B, Table S5) and seven of the top 50 genes from *C. ljungdahlii* were annotated as pyrimidine biosynthesis proteins (Fig. 2A, Table S6). This would appear to indicate that, especially early in coculture, both coculture partners derive a growth benefit from the other’s presence. It is clear why *C. ljungdahlii* growth would improve in coculture; it receives CO_2_, H_2_, and (as suggested in the previous section) amino acids from *C. acetobutylicum.* Although *C. ljungdahlii* does not directly provide any growth substrates for *C. acetobutylicum,* we hypothesized that “local” (that is, at the microscale cell-to-cell interaction level, as suggested by the Type II comparison) consumption of H_2_ by *C. ljungdahlii* would relieve feedback inhibition of *C. acetobutylicum’s* hydrogenase (29). In *C. acetobutylicum,* during the acidogenic growth phase (and especially early acetogenic phase (30, 31)), a high fraction of electrons from glycolysis are used to produce H_2_ (32), thus enabling faster cell growth and consumption of glucose. It is well established that different forms of in situ H_2_ removal, including via coculture with an H_2_-metabolizing organism, can improve the growth and H_2_ production rate of carbohydrate fermenting *Clostridium* species (33) (34). Based on this hypothesis, we measured the rate of glucose consumption in the first few hours (early to mid-exponential growth, per the guidance from (30, 31)) of a *C. acetobutylicum* monoculture under either a pure N_2_ headspace or an 80% H_2_, 20% CO_2_ headspace compared to the rate in coculture with *C. ljungdahlii*. We observed that the rate of glucose consumption in the coculture is as high or higher than the N_2_ control, and higher than the 80% H_2_ control, at each timepoint examined in the first six hours in all three biological replicates (Fig. S3). The effect was small, but was present in all three biological replicates, consistent with the hypothesis that H_2_ removal at the local level due to its utilization by *C. ljungdahlii*, would benefit *C. acetobutylicum* by reducing the feedback inhibition of its hydrogenase.

### Cell Motility Genes in C. ljungdahlii and C. acetobutylicum

While the mechanism that leads to heterologous, interspecies, cell-wall and eventually cell-membrane fusion remains unchartered territory in prokaryotic biology, mammalian-cell fusion has been widely studied (35, 36): the process involves several essential steps: chemotaxis and migration, membrane adhesion, membrane fusion, and post-fusion resetting. Chemotactic attraction and forces are essential for mammalian cell fusion (36) to enable close cell contact by overcoming the repelling forces due to cell’s surface charges. Based on fusion process of mammalian cells and Transmission Electron Microscopy (TEM) images (2), we hypothesized that the fusion process in our system involves analogous steps, starting with chemotaxis and cell migration (“swimming”). Given *C. ljungdahlii*’s absolute need for CO_2_ and H_2_ for survival, one would assume that *C. ljungdahliij* would be chemotactically attracted to CO_2_ and H_2_ and thus to *C. acetobutylicum* that produces them during glucose catabolism. As there are no published chemotactic (“swim”) assays towards gaseous attractants, we developed a novel swim assay (37) to show that *C. ljungdahlii* is indeed attracted to a 80% H_2_/ 20% CO_2_ gas mix or to CO_2_ alone. Here, we wanted to test if our transcriptional data would support this hypothesis. One can distinguish two scales of *C. ljungdahlii* “swimming”: in mixed, high-density cocultures with *C. acetobutylicum*, *C. ljungdahlii* would swim short distances to reach neighboring *C. acetobutylicum* cells, the source of CO_2_ and H_2_. However, when *C. ljungdahlii* is grown in monoculture with an 80% H_2_/ 20% CO_2_ atmosphere in serum bottles (Fig. 1A), *C. ljungdahlii* cells need to travel longer distances, on average, to move towards the liquid surface which has the highest gas (H_2_+CO_2_) concentrations (partial pressures). One might expect then that the motility genes would be expressed higher in *C. ljungdahlii* monoculture vs in *C. acetobutylicum*-*C. ljungdahlii* cocultures. Indeed, in the Type I experiment, at 4 and 11 hours, *C. ljungdahlii* cell motility genes were largely expressed lower in coculture compared to monoculture (Table S10). The flagellin gene (CLJU_c09630) coding for the flagellin protein, which polymerizes to form the flagella was significantly downregulated in coculture. Almost all of the genes of the large flagellar operon (CLJU_c10180-10450) were downregulated at the 4-hr timepoint or both the 4 and 11-hour timepoints, including the *fliG* gene, which codes for the flagellar motor switch protein FliG. The smaller operon (CLJU_c09470-09630) contains two genes (*fliM, fliN*) that were upregulated at 4 and 11 hours. FliM and FliN are partners of the FliG protein, and in this sense it is not clear why they display a different expression pattern. The two *mot* genes (CLJU_c01450-01460), coding for proteins of the flagellar motor, were downregulated at the 4hr timepoint. As hypothesized then, for *C. ljungdahlii*, the clear trend appears to be lower expression of the flagellar complex early in coculture for the Type I experiments. This behavior of cell motility genes was not observed in Type II experiments (Table S10) because of the lack of mixing. In the transwell system, *C. ljungdahlii* cells in the top well can simply settle to the bottom of the well to access the CO_2_ and H_2_ produced by *C. acetobutylicum* in the bottom well, as opposed to the mixed monoculture in which they must fight gravity and mixing to access the CO_2_ and H_2_ available at the gas-liquid interface. This settling phenomenon was experimentally observed; in all separated transwell experiments *C. ljungdahlii* cells formed a defined layer at the bottom of the transwell.

In *C. acetobutylicum,* in the Type I experiments, there was no differential expression of flagellar complex genes or other genes associated with cell motility. However, in *C. acetobutylicum,* sixteen different flagellar genes (CA_C2143, 2147, 2156-2164) were expressed higher in unseparated vs separated coculture, largely at 4 hours (Table S11). This finding further supports the conclusion that *C. acetobutylicum* benefits from in situ H_2_ removal by *C. ljungdahlii.* These results suggest that, similar to how *C. ljungdahlii* swims towards higher concentrations of CO_2_ and H_2_, *C. acetobutylicum* may swims towards lower concentrations of H_2_ to minimize repression of its hydrogenase protein and enable faster consumption of glucose, as described in a previous section. Excitingly, these findings suggest that chemotactic attraction in the coculture is likely not just one way (*C. ljungdahlii* swimming towards *C. acetobutylicum*) but a two-way process, suggesting that both organisms may play an active role in the hypothesized first step of the fusion process: chemotaxis and migration.

### Key Wood-Ljungdahl Pathway (WLP) and Autotrophic Energy Metabolism Genes are Either Downregulated or Unaffected in *C. ljungdahlii* in Coculture compared to *C. ljungdahlii* monocultures under a H_2_/CO_2_ atmosphere

Several studies have reported gene-expression patterns between *C. ljungdahlii* grown heterotrophically and autotrophically (38, 39). Here we focused on genes impacted by the coculture with *C. acetobutylicum*. For this reason, we cultured the *C. ljungdahlii* monoculture controls under a 20 psig atmosphere with 80% H_2_ and 20% CO_2_. Because in both monoculture and coculture *C. ljungdahlii* has access to a good supply of CO_2_ and H_2_, we hypothesized that there will be no differential expression or, possibly, downregulation of WLP and associated genes in the coculture. WLP-gene downregulation may be based on the feeding off the arginine produced by *C. acetobutylicum*, as discussed above. Downregulation of the WLP genes would also be supported by the experimental evidence that *C. ljungdahlii*, beyond feeding on CO_2_ and H_2_ and amino acids produced by *C. acetobutylicum*, may partially feed off of *C. acetobutylicum*’s intracellular metabolites and reduced electron carriers, via the large scale exchange of cytoplasmic material during heterologous cell fusion (2, 8), thus decreasing its reliance on autotrophic metabolism.

Indeed, in the Type I experiments, we observed that the central WLP genes either were not differentially expressed or were downregulated under coculture conditions (Table S12): several subunits of cooS1 (CLJU_c09090-09110), one of *C. ljungdahlii*’s three CO dehydrogenases, one of its two formate dehydrogenase genes (CLJU_c08930), and *C. ljungdahlii’s* primary uptake hydrogenase (CLJU_c07070) were downregulated at 4 and 11 hours. This is notable because the CO dehydrogenase governs the conversion of CO_2_ to CO required for the first step of the western branch of the WLP, and formate dehydrogenase converts CO_2_ to formate in the first step of the eastern WLP branch. The hydrogenase protein is responsible for H_2_ utilization and generating reducing equivalents for the WLP when *C. ljungdahlii* is grown in the presence of H_2_. Taken together, the downregulation of genes involved in the first steps of both branches of the WLP and H_2_ uptake suggest that, early in coculture, *C. ljungdahlii* autotrophic carbon fixation is downregulated (Fig. 7A). A similar trend was observed for the Type II experiments. For the Type II, the three subunits of cooS1 were not downregulated, but there was downregulation of the contiguous ferredoxin gene (CLJU_c07060) associated with the H_2_ uptake hydrogenase at both 4 and 11 hour (only 11 hour in the Type I experiment).

**Figure 7:**
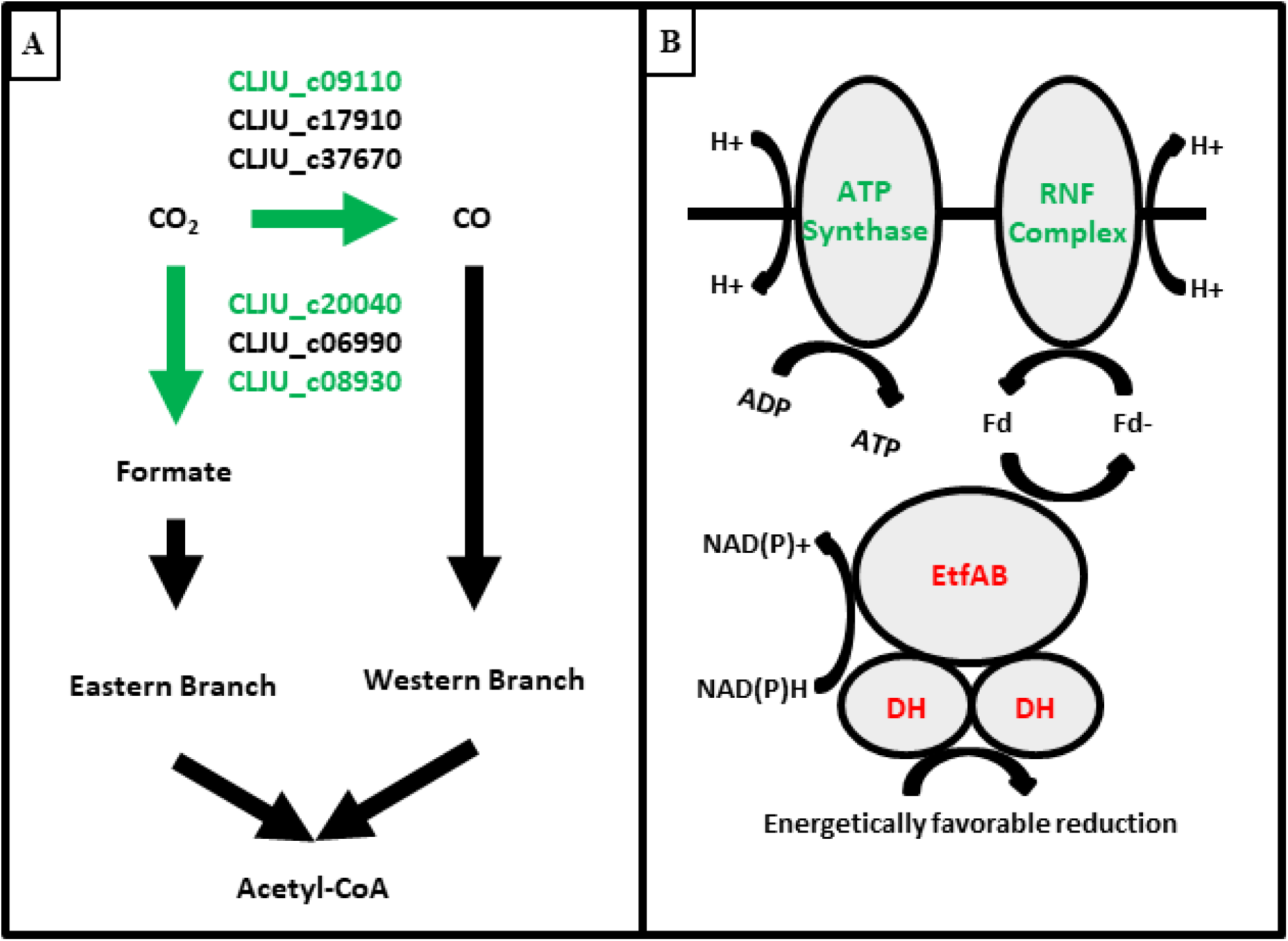
A) Schematic of Wood-Ljungdahl Pathway annotated with C. ljungdahlii accession numbers of the gene products (enzymes) that catalyze key reaction steps. B) Schematic depicting the metabolic function of ATP synthase and EtfAB enzymes. Red or green lettering of accession numbers or protein complexes indicate that genes coding for those proteins are up or down regulated at one or more timepoints in either the Type I or Type II experiments.

In addition to WLP genes, several key genes involved in the coupling of the WLP and energy production in *C. ljungdahlii* were also downregulated (Table S13). In the Type I experiments two subunits of the Rnf complex were downregulated at 4 hours (CLJU_c11370-11380), and those two subunits in addition to four other contiguous (CLJU_c11360, 11390-11410) Rnf subunit genes were downregulated at 11 hours. Three subunits of *C. ljungdahlii*’s ATP synthase, which works in concert with the Rnf complex to generate ATP, were also downregulated at 11 hours. Interestingly, two of *C. ljungdahlii*’s three annotated etfAB operons were strongly upregulated at one or both of the 4- and 11-hour timepoints (CLJU_c13880-13900, 21570-21590). These operons consist of three proteins: the two subunits of an electron transfer flavoprotein complex and a dehydrogenase that performs coupled redox reactions with the etfAB complex in order to conserve energy or to enable otherwise energetically unfavorable reductions (40). The dehydrogenases associated with the upregulated etfAB genes are annotated as lactate (CLJU_c21570) and glycolate (CLJU_c13900) dehydrogenases. The strong upregulation of these etfAB-dehydrogenase operons in coculture may indicate that the *C. ljungdahlii* enzymes can use metabolites from *C. acetobutylicum* to carry out energetically favorable redox reactions to support its own metabolism. Downregulation of subunits of key autotrophic metabolism genes, such as the Rnf and ATPase synthase complexes, combined with the upregulation of the etfAB operons (Fig. 7B), further support the hypothesis that *C. ljungdahlii* can afford to downregulate certain genes associated with autotrophic energy production because of metabolites and electrons it obtains through diffusion as well as the transfer of cytoplasmic material from *C. acetobutylicum* through interspecies fusion (2) in coculture.

## CONCLUSIONS

The gene expression data paint a picture in which *C. ljungdahlii* gene expression is heavily modified based on nutrients it is able to receive from *C. acetobutylicum*, as well as the increased ease of obtaining nutrients via direct exchange in addition to CO_2_, H_2_ and amino acids through the liquid medium. The pattern of amino acid biosynthesis differential expression, combined with amino acid supernatant profiles, suggest that *C. ljungdahii* may obtain several key amino acids directly from *C. acetobutylicum*. These amino acids may then be used for protein synthesis, ATP generation or conservation via catabolic reactions such as the arginine deiminase pathway, histidine catabolism to glutamate, and peptidoglycan biosynthesis. Despite the increased amount of CO_2_ and H_2_ available to *C. ljungdahii* in coculture with *C. acetobutylicum*, the genes of the WLP and *C. ljungdahii’s* autotrophic energy metabolism were downregulated. This suggests that, in coculture, *C. ljungdahii* has access to sources of nutrients (notably amino acids produced by *C. acetobutylicum*), and, thus, the enzymes of the WLP and the associated machinery for ATP production can be expressed at lower levels. In *C. acetobutylicum*, the pattern of ribosomal protein expression combined with the known impact of H_2_ partial pressure (inhibition of its hydrogenase) and the simple experiments of Fig. S3 suggest that *C. acetobutylicum* derives a growth benefit from its coculture with *C. ljungdahii,* thus suggesting that this is mutualistic syntrophy. Finally, differential regulation of cell motility genes suggests that chemotactic cell movement is affected by the coculture, as we would expect based on previous results showing chemotactic attraction (37) of *C. ljungdahii* towards CO_2_ + H_2_ (and by inference to *C. acetobutylicum* that produces the two gases) as first step leading to heterologous cell fusion (2, 8) between the two organisms.

## MATERIALS AND METHODS

### Microorganisms and culture media

Monocultures of *C. acetobutylicum* ATCC 824 and *C. ljungdahlii* DSM 13528 as well as cocultures combining the species were grown for RNA extraction. For *C. acetobutylicum*, a frozen stock was streaked onto a 2xYTG agar plate (41) and left at 37 C in an anaerobic incubator for 3 days to allow for colony growth and spore formation. All subsequent culturing steps were performed in Turbo CGM growth medium, a complex medium formulation (containing 5 g/L yeast extract) designed to simultaneously support the growth of both *C. acetobutylicum* and *C. ljungdahlii* (8). After incubation, a single colony was inoculated into a test tube containing 10 mL of Turbo CGM growth medium, heat shocked at 80 C to kill non-sporulated cells, and grown statically in an anaerobic incubator at 37 C for 16-24 hours. The resulting pre-culture was used to inoculate 30 mL of Turbo CGM growth medium to an OD_600_ of 0.1. This culture was grown statically in the anaerobic incubator at 37 C until it reached exponential phase (OD_600_ between 0.4-0.6). For both the coculture and monoculture controls, Turbo CGM was inoculated with 17% of this exponential pre-culture to a final volume of 60 mL. *C. acetobutylicum* monocultures and cocultures were grown in 100 mL screw cap bottles and grown statically at 37 C in an anaerobic incubator. The screw caps were loosened so that gases produced by the culture could escape to avoid building potentially explosive gas pressure.

For *C. ljungdahlii*, a frozen stock was inoculated into 30 mL of Turbo CGM growth medium without glucose in a 160 mL serum bottle which had been pre-flushed for 2 min with an 80% H_2_, 20% CO_2_ gas mix. The serum bottle was pressurized to 20 psig with the 80/20 gas mixture and grown for 24 hours at 37 C on a rotating platform at 90 rpm. This pre-culture was used to inoculate pre-flushed serum bottles each containing 75 mL of Turbo CGM without glucose to an OD_600_ of 0.1. These pre-cultures were grown to exponential phase (OD_600_ of 0.4-0.6) and used to inoculate the cocultures and *C. ljungdahlii* monocultures. For coculture preparation, a volume of exponential *C. ljungdahlii* pre-culture corresponding to an R (ratio of *C. ljungdahlii* to *C. acetobutylicum* cells based on OD_600_) = 5 (8) was centrifuged at 5000 rpm, 4 C for 10 min, resuspended in 1mL of Turbo CGM, and added to the coculture with a final volume of 60 mL. For monoculture preparation, 1.5 times this volume was centrifuged at 5000 rpm, 4 C for 10 min, resuspended in 1mL of Turbo CGM without glucose, and used to inoculate Turbo CGM without glucose to a final volume of 90 mL. *C. ljungdahlii* monocultures were grown in sealed 160 mL serum bottles which had been pre-flushed for 2 min with an 80% H_2_, 20% CO_2_ gas mix. After culture addition, the serum bottles were pressurized to 20 psig with the 80/20 gas mixture and grown at 37 C and 90 rpm. Each condition was prepared in biological triplicates.

### Transwell cultures

A transwell system (Corning, Prod. No. 7910) was used to prepare co-cultures for RNA extraction in which *C. acetobutylicum* and *C. ljungdahlii* were separated by a permeable membrane (0.4 µm pore size). Each transwell plate contained 6 wells and 5 membrane inserts (the membrane was removed from one of the wells on each plate to allow culture mixing). Each of these wells had a total volume of 8.0 mL (4 mL in the bottom of the well and 4 mL in the membrane insert). To prepare membrane separated cultures, *C. acetobutylicum* was added to fresh Turbo CGM at 34% inoculum to a final volume of 4 mL and added to the bottom of the well. A volume of *C. ljungdahlii* corresponding to an R value of 5 was centrifuged at 5000 rpm, 4 C for 10 min, resuspended in 4 mL of Turbo CGM, and added to the top of the well inside the membrane insert or directly to the well if the insert was omitted. Three transwell plates were prepared, one each for the 2, 4, and 11-hour timepoints.

### RNA Sample Preparation

Cell samples for RNA extraction were taken 4, 11, and 22 hours after culture inoculation for all bottle cultures. Cell samples for RNA extraction were taken from the transwell cultures at 2, 4, and 11 hours after culture inoculation. For the wells containing membrane inserts, the *C. ljungdahlii* suspensions were harvested from each of the five well plates containing the membrane inserts and pooled. After harvesting *C. ljungdahlii*, the membrane inserts were removed, and the *C. acetobutylicum* suspensions were harvested from the bottom of the wells and pooled. All cell samples for RNA extraction were centrifuged at 5000 rpm, 4 C for 10 min. The supernatant was decanted, and the pellets were resuspended in a 2:1 solution of PBS and Qiagen RNAprotect Bacterial Reagent and left to incubate at room temperature for 10 min. After incubation, the cells were re-pelleted by centrifugation at 5000 rpm, 4 C for 10 min. The supernatant was decanted, and the pellets were rinsed with 1 mL RNase-free SET buffer (25% sucrose, 50 mM EDTA, 50 mM Tris, 20 mM DTT, heat treated at 60 C for 10 min). Following that, the pellet was resuspended in 500 uL of RNase free SET buffer with 20 mg/mL lysozyme and 60 mAU/mL proteinase K and left to incubate for 6 min at room temperature. Cell solutions were then transferred to 2 mL RNase free tubes containing 40 mg of RNase-free glass beads (bead diameter ≤ 106 µm) and 1 mL of ice cold Qiazol reagent and placed on a Vortex-Genie 2 with an epitube adapter. Samples were vortexed at maximum speed for 4 min and then immediately stored at −80 C.

For RNA preparation, the cell-Qiazol suspensions were allowed to thaw on ice. 500 ul of lysate was combined with 500 ul of fresh ice-cold Qiazol and 200 ul of ice-cold chloroform was added. Each solution was mixed thoroughly with 4-5 inversions, left to incubate at room temperature for 10 minutes, and centrifuged at 13,000 rcf, 4 C for 15 min. The aqueous phase (approximately 700 ul per sample) was transferred to a fresh epitube and combined with 1.3 mL of ice-cold 100% ethanol. This solution was mixed thoroughly via inversion and transferred to a Qiagen RNeasy spin column. RNA was bound, cleaned, and treated with DNase according to the kit protocol and optional on-column DNase treatment. The final product was eluted in 100 ul of nuclease free water.

RNA yield and purity were quantified with a Denovix DS-11 Spectrophotometer and RNA integrity was assessed using the High Sensitivity RNA Analysis mode on an Agilent Fragment Analyzer. Based on the eluate concentration, a volume of each sample solution corresponding to 1 ug of total RNA was purified and concentrated to the required library prep input concentration using the Zymo Clean and Concentrator kit according to the manufacturer protocol. Samples were eluted in 8.5 ul of nuclease free water and stored at −20 C until RNAseq library preparation. Leftover RNA samples were precipitated via the addition of 0.1 volumes 3 M sodium acetate and 2.5 volumes ice-cold 100% EtOH and stored at −20 C.

### RNAseq Library Preparation, Sequencing & Data Analysis

Ribosomal RNA depletion and RNAseq library preparation were performed concurrently using the Zymo-Seq RiboFree Total RNA Library Kit according to the manufacturer protocol following the recommended modifications for degraded RNA samples. Library yields were quantified on a Qubit fluorometer using the dsDNA High Sensitivity assay, and library size distribution was quantified using the High Sensitivity NGS mode on an Agilent Fragment Analyzer. RNAseq libraries were pooled for sequencing, with libraries representing coculture samples added at a 4x higher concentration than the monoculture libraries. Pooled libraries from the bottle cocultures were sequenced on one lane of an Illumina NextSeq 550 using a 150 cycle kit to generate 2×76 bp paired end reads using the high output option. Pooled libraries from the transwell cocultures were sequenced on one lane of an Illumina NextSeq 2000 using a 200 cycle kit to generate 2×100 bp paired end reads using the high output option.

Paired-end sequencing for the bottle (Type I) experiments produced 714,191,023 pairs of 76 bp reads; paired-end sequencing for the transwell (Type II) experiments produced 714,191,023 pairs of 100 bp reads. Sequencing files were uploaded to the Galaxy bioinformatics platform for adapter trimming and quality control. Sequencing adapters were removed from the reads using the provided Zymo-Seq RiboFree Total RNA Library Kit adapter sequences and Trim Galore! software accessed through the Galaxy bioinformatics platform. Read quality control was performed using FastQC software accessed through the Galaxy bioinformatics platform. Read alignment, differential expression analysis, and operon detection were performed with the open source prokaryotic RNAseq software Rockhopper (42, 43) using default parameter settings. *C. acetobutylicum* monoculture samples were aligned to the *C. acetobutylicum* ATCC 824 genome and pSOL1 megaplasmid, *C. ljungdahlii* monoculture samples were aligned to the *C. ljungdahlii* DSM 13528 genome, and coculture samples were aligned against all three. Genes with q-value < 0.05 and log-fold change > |1| were considered differentially expressed (44). All sequencing reads generated in this study are deposited in NCBI’s sequence reads archive under the BioProject PRJNA1167669 containing 54 BioSamples (SAMN44010615-SAMN44010668).

### Metabolite assays

All bottle co-cultures and mono-cultures were sampled at 4, 11, 22, and 50 hours. All transwell cultures were sampled at 2, 4, and 11 hours when the plates were harvested for RNA extraction. High pressure liquid chromatography (HPLC) was used to quantify supernatant glucose, fructose, and solvent concentrations as previously reported (45–47). For amino acid quantification, cell supernatants were diluted 20-fold and analyzed on an Agilent 1260 Infinity II HPLC equipped with an Agilent Poroshell HPH-C18 column. Mobile Phase A was a solution of 10 mM dibasic sodium phosphate, 10 mM sodium borate, pH adjusted to 8.2 with hydrochloric acid. Mobile Phase B was a solution of 45% methanol, 45% acetonitrile, and 10% water. All samples were ran using an 18 min gradient elution with a flowrate of 1.5 ml/min. Results were compared against a compared against a calibrate curve of amino acid standards purchased from Agilent (Product numbers 5061-3330, 5061-3331, 5061-3332, 5061-3333, 5061-3334). For ammonia quantification, cell supernatants were diluted 5-fold and analyzed on a YSI 2950D Biochemistry Analyzer using the installed ammonium biosensor module.

## SUPPLEMENTAL MATERIAL

**Supplemental Figures (Word document)**

Figures S1-S3

**Tables S1-S4 Excel, multiple sheets)**

Complete summaries of the differential and absolute gene expression data for both the Type I and II RNAseq experiments.

**Tables S5-S13 (Excel, multiple sheets)**

Supplemental gene expression heatmaps

**Table S14-S15 (Excel, multiple sheets)**

KEGG Pathway Analysis for both the Type I and II RNAseq experiments.

**Supplementary Document 1 for the analysis of the data presented in Figure S2 and Table S3**

## ACKNOWLEDGEMENTS

This work was supported by an ARPA-E project under contract AR0001505. N.B.W. was supported in part by a U.S. Department of Education GAANN Fellowship under grant P200A210065.

We thank Dr. Bruce Kingham and Mark Shaw of the University of Delaware Sequencing and Genotyping Center for assistance with Illumina RNA sequencing.

E.T.P. conceived the project. E.T.P. and N.B.W. designed the experiments. N.B.W. performed all experiments. E.T.P. and N.B.W. analyzed the data and wrote the manuscript.

## Supplementary Text

**Supplementary Text Document 1 for the analysis of the data presented in Figure S2 and Table S3**

**Timepoint Overlap and KEGG Pathway Analysis of Differential Gene Expression in Type I and II Experiments**

All differentially expressed genes were classified by pathway according to the KEGG database and analyzed via simple functional enrichment using the hypergeometric test (48) to determine which KEGG pathways were statistically overrepresented in the differentially expressed gene list for each species, timepoint, and subsection (Table S14, S15). All genes differentially expressed in *C. acetobutylicum* and *C. ljungdahlii* at a given timepoint in one or both RNAseq comparisons, as well as the subfractions of genes that were differentially expressed at multiple timepoints are illustrated by the Venn diagrams of Figs. S2A and S2B. The major conclusion from the KEGG pathway analysis was that coculture significantly impacts gene expression of amino acid metabolism genes in both organisms in both the Type I and II experiments, especially biosynthesis of arginine, histidine, and tryptophan. Many other KEGG pathways were differentially regulated in each individual organism at different timepoints, but the impact on amino acid metabolism was seen in both organisms across all timepoints in both the Type I and II experiments.

## Supplementary Figures

**Figure S1:**
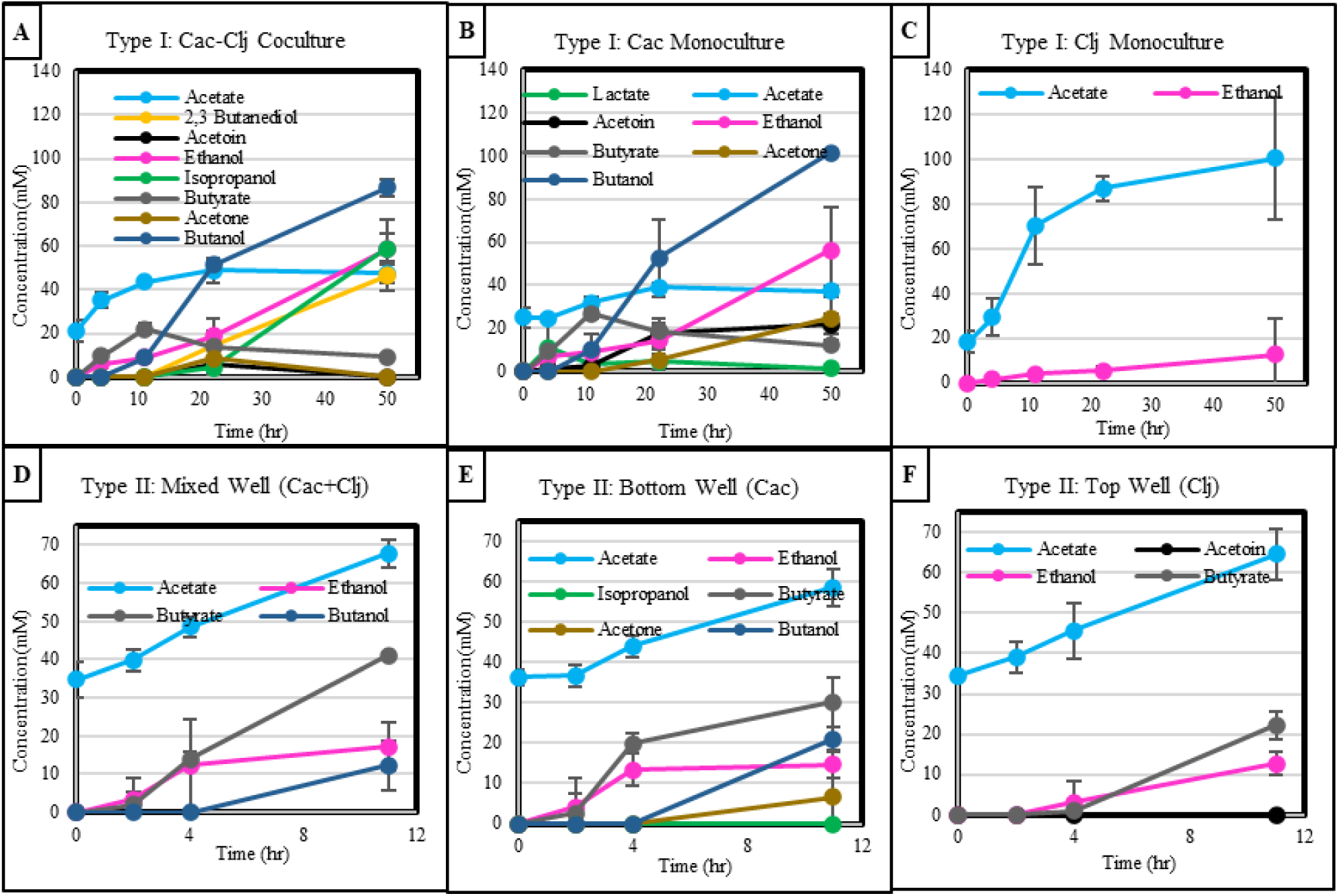
Temporal metabolite profiles. A) Type I C. acetobutylicum (Cac)- C. ljungdahlii (Clj) cocultures. B) Type I C. acetobutylicum monoculture. C) Type I C. ljungdahlii (monoculture. D) Type I unseparated C. acetobutylicum- C. ljungdahlii coculture. E) Type II bottom well (C. acetobutylicum) culture. F) Type II top well (C. ljungdahlii) culture.

**Figure S2:**
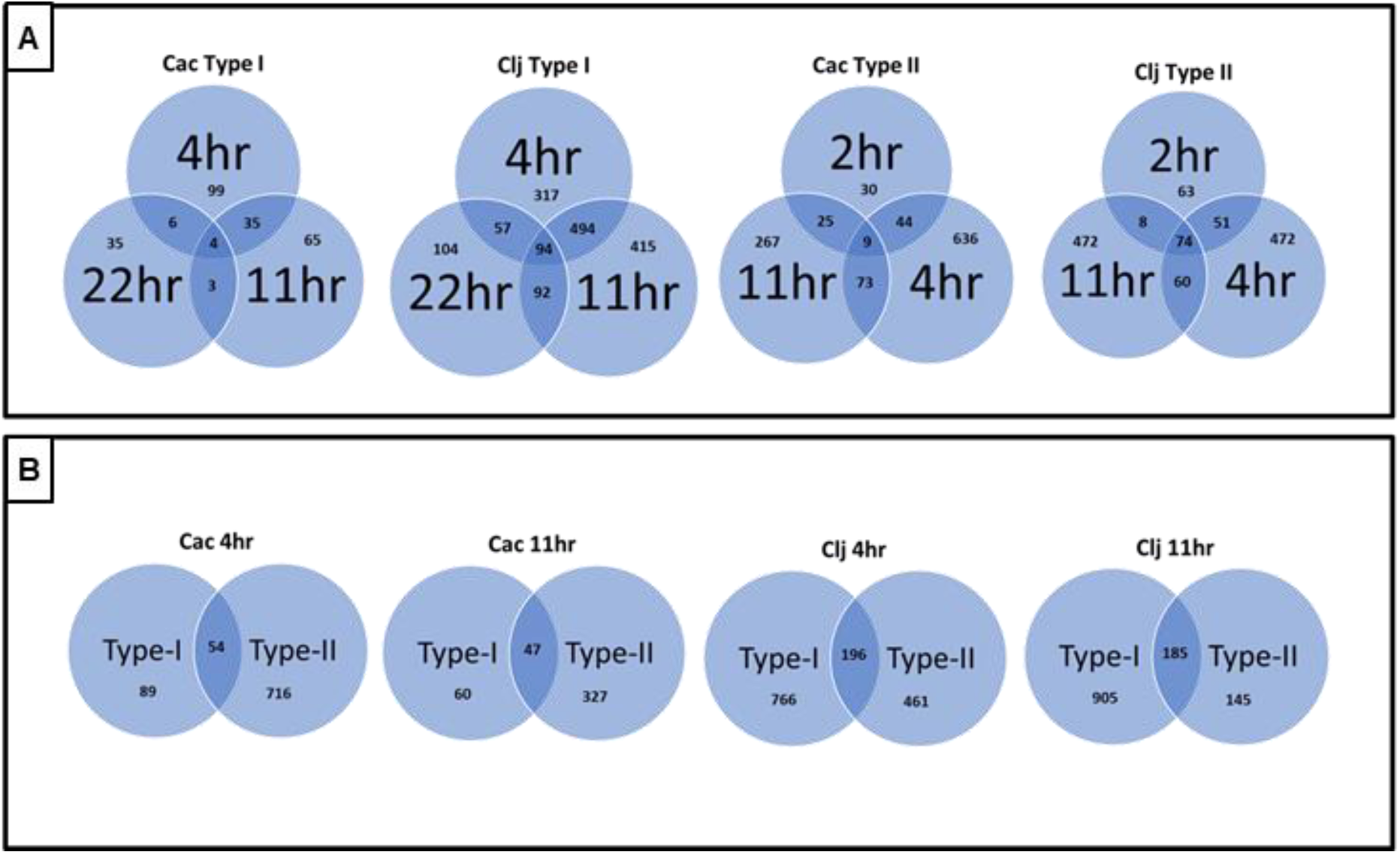
A) Venn diagrams of differentially expressed genes at one or more timepoints for C. acetobutylicum (Cac) and C. ljungdahlii (Clj) in Type I and Type II experiments. B) Venn diagrams of differentially expressed genes in either the Type or Type II or both experiments at each timepoint in both C. acetobutylicum and C. ljungdahlii. Venn diagrams represent global picture of gene expression as described in Supplementary Document 1.

**Figure S3:**
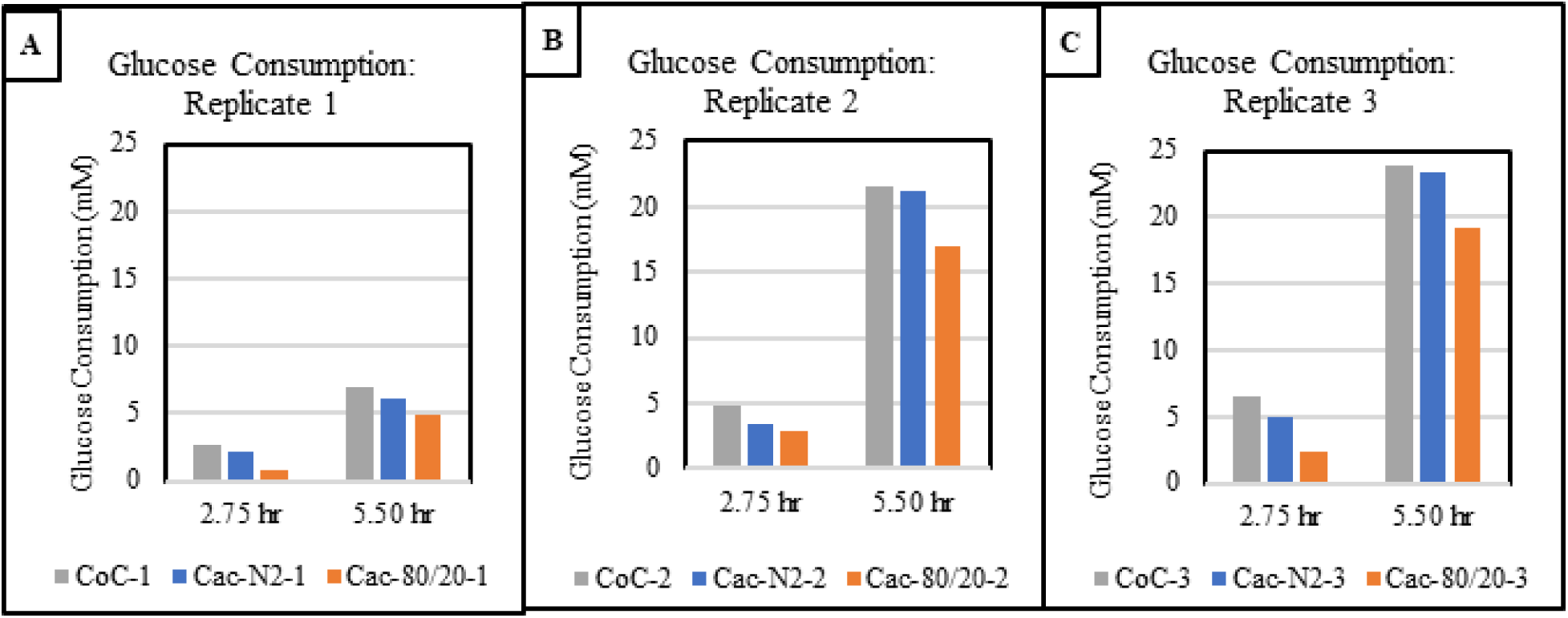
Cumulative glucose consumption at each timepoint for C. acetobutylicum (Cac)- C. ljungdahlii (Clj) coculture (CoC), C. acetobutylicum monoculture with N_2_ headspace (Cac-N2), and C. acetobutylicum monoculture with 80% H_2_ / 20% CO_2_ headspace (Cac-80/20). Three biological replicates (A, B, C) shown individually.

